# Isolation and characterization of bovine herpes virus 5 (BHV5) from cattle in India

**DOI:** 10.1101/2020.01.24.917880

**Authors:** Naveen Kumar, Yogesh Chander, Thachamvally Riyesh, Nitin Khandelwal, Ram Kumar, Harish Kumar, Bhupendra N. Tripathi, Sanjay Barua

## Abstract

Bovine herpesvirus 1 (BHV1) and 5 (BHV5) are genetically and antigenically related alphaherpesviruses. Infection with one virus induces protective immunity against the other. However, disease associated with BHV1 and BHV5 varies significantly; whereas BHV1 infection is usually associated with rhinotracheitis and abortion, BHV5 causes encephalitis in cattle. BHV5 outbreaks are sporadic and mainly restricted to the South American countries. We report BHV5 infection for the first time from aborted cattle in India. Based on the characteristic cytopathic effects in MDBK cells, amplification of the viral genome in PCR, differential PCR for BHV1/BHV5, nucleotide sequencing and restriction endonuclease patterns, identity of the virus was confirmed as BHV5 subtype A. Serum samples from the aborted cattle strongly neutralized both BHV1 and BHV5 suggesting an active viral infection in the herd. Upon *UL27, UL44* and *UL54* gene-based sequence and phylogenetic analysis, the isolated virus clustered with BHV5 strains and showed highest similarity with the Brazilian BHV5 strains.

**Author Summary:** BHV5 causes fatal meningoencephalitis that may result in a mortality rate of up to 100% in cattle. BHV5 is rarely associated with abortion and its distribution is restricted to South American countries. Only a few cases of this disease have been reported from other countries. For the first time, we provide a strong evidence of BHV5 infection from aborted cattle in India. The finding may necessitate inclusion of BHV5 test protocol in testing of semen for sexually transmitted diseases. Also, the isolated virus would be useful for developing diagnostic, prophylactic and therapeutic agents to combat BHV5 disease in the country.

## Introduction

Bovine herpesvirus types 1 (BHV1) and 5 (BHV5) belong to the family *Herpesviridae*, subfamily *Alphaherpesvirinae* and genus varicellovirus [1]. They are genetically and antigenically closely related and share ~85% nucleotide identity. BHV1 is prevalent globally and causes infectious bovine rhinotracheitis (IBR) in cattle which is characterized by conjunctivitis, rhinotracheitis, balanopostitis, infectious pustular vulvovaginitis, enteritis, abortion and sometimes encephalitis. BHV1 infection results in considerable economic losses due to decreased milk production, weight loss and abortions.

BHV5 usually infects young calves and mortality can causes up to 100% [2]. BHV5 mainly causes fatal meningoencephalitis in cattle and establishes latency in trigeminal ganglion. The virus is excreted in nasal, ocular and genital secretions upon reactivation (under stress). The clinical signs in affected cattle include depression, anorexia, weakness followed by neurological signs, such as incoordination, muscular tremor, blindness, circling, recumbency, head pressing, convulsions, paddling and death [3]. Occasionally, BHV5 has been shown to be associated with the reproductive disorders [4].

Herpesvirus associated bovine meningoencephalitis was first time reported in 1962 in Australia. Based on the virion morphology, cytopathic effect in cell culture and antigenic properties, the isolated virus was initially considered to be a neuropathogenic variant of BHV1, called bovine encephalitis herpesvirus (BEHV) [5] or BHV1 subtype 3 [1]. However, later on, based on the restriction site mapping of viral DNA and cross reactivity with monoclonal antibodies, BEHV was found to be a distinct strain with different genomic and antigenic properties. Thus, in 1992, International Committee on Taxonomy of Viruses recognised BEHV as a distinct virus species, namely BHV5 [6].

The prevalence of BHV5 is not precisely known because the available serologic tests do not discriminate antibodies against BHV1- and BHV5. Naturally occurring or vaccine-induced BHV1 antibodies confer cross protection against BHV5, a possible reason of non-occurrence of BHV5-associated disease in BHV1 endemic areas [7,8].

BHV5 is endemic in South American countries-Argentina [9], Uruguay [10] and Brazil [11]. Only a few cases of the disease have been reported in Australia [12], Hungary [13], Iran [14], Canada [15] and United States [16]. BHV5 has not been reported in India. We isolated and characterized BHV5 for the first time from aborted cattle in India.

## Materials and Methods

### Ethics Statement

The study involves collection of biological specimens from cattle (field animals). Vaginal swabs and blood samples (3 ml each) were collected from three aborted and three apparently healthy cattle as per the standard practices without using anaesthesia. ICAR-National Research Centre on Equines, Hisar (India) granted the permission to collect the biological specimens. A due consent was also taken from the farmer (animal owner) before collection of the specimens.

### Cells and virus

Madin-Darby bovine kidney (MDBK) cells were grown in Dulbecco’s Modified Eagle’s Medium (DMEM) supplemented with antibiotics and 10% fetal calf serum. Reference BHV1 (VTCCAVA14) was available in our repository at National Centre for Veterinary Type Cultures (NCVTC), Hisar, India which has been described elsewhere [17,18].

### Clinical specimens

The samples originated from an organized cattle farm located near Bhilwara, Rajasthan, India (25.3407° N, 74.6313° E). Out of a total of 68 animals, 61 (including 2 bulls) and 7 (including one bull) belonged to Gir and Tharpakar breeds, respectively. First evidence of abortion in the farm was noticed about 2 years prior to the sampling. Over 25 abortions had already occurred. Three cattle with a recent history of abortion (3 -18 days prior to sampling) as well as 3 apparently healthy animals without any history of abortion were considered for sampling. Abortions occurred between 4-9^th^ months of pregnancy without any specific pattern. No involvement of nervous system was recorded in any of the aborted cows. The farmer practiced natural service in the farm and never performed artificial insemination for breeding purpose. Besides six-monthly vaccinations against foot-and-mouth disease (FMD), the farmer also employed RB51 calfhood vaccination against *Brucella*. Bulls for breeding were purchased from Surendranagar, Gujarat, a nearby state (India).

Vaginal swabs were collected in Minimum Essential Medium (MEM, transport medium) and transported on ice. The biological specimens were processed as per the standard procedures described elsewhere [19]. An aliquot of 500 μl suspension of vaginal swab was used for bacteriological examinations. The remaining sample was filtered through 0.45 μM syringe filters and used for various virological assays. Serum/blood samples (3 ml from each animal) were also collected from aborted as well as healthy animals. A due consent was taken from the animal owner for collection of the biological specimens.

### Identification of the agent(s)

Initially, the DNA was extracted from vaginal swabs by DNeasy Blood & Tissue Kits (Qiagen, Valencia, CA, USA) and subjected to amplification of the BHV1, *Campylobacter spp, Brucella spp, Leptospiran spp*, *Listeria spp* and *Trichomonas vaginalis* genome, the agents which are commonly associated with abortion in cattle. Primers, annealing temperatures and extension times for amplification of these agents are given in **Table 1.** For PCR amplification, each reaction tube of 20 μl contained 10 μl of Q5 High-Fidelity 2× Master Mix (New England BioLabs Inc.), 20 pmol of forward and reverse primer, and 5 μl of DNA (template). Thermocycler conditions included: a denaturation step of 5 min at 98°C followed by 35 cycles of amplification [(30 sec at 98°C, 30 s at 58-65°C **(Table 1)** and 40-90s **(Table 1)** at 72°C], and a final extension step at 72°C for 10 min. The PCR products were separated in a 1% agarose gel.

**Table 1:**
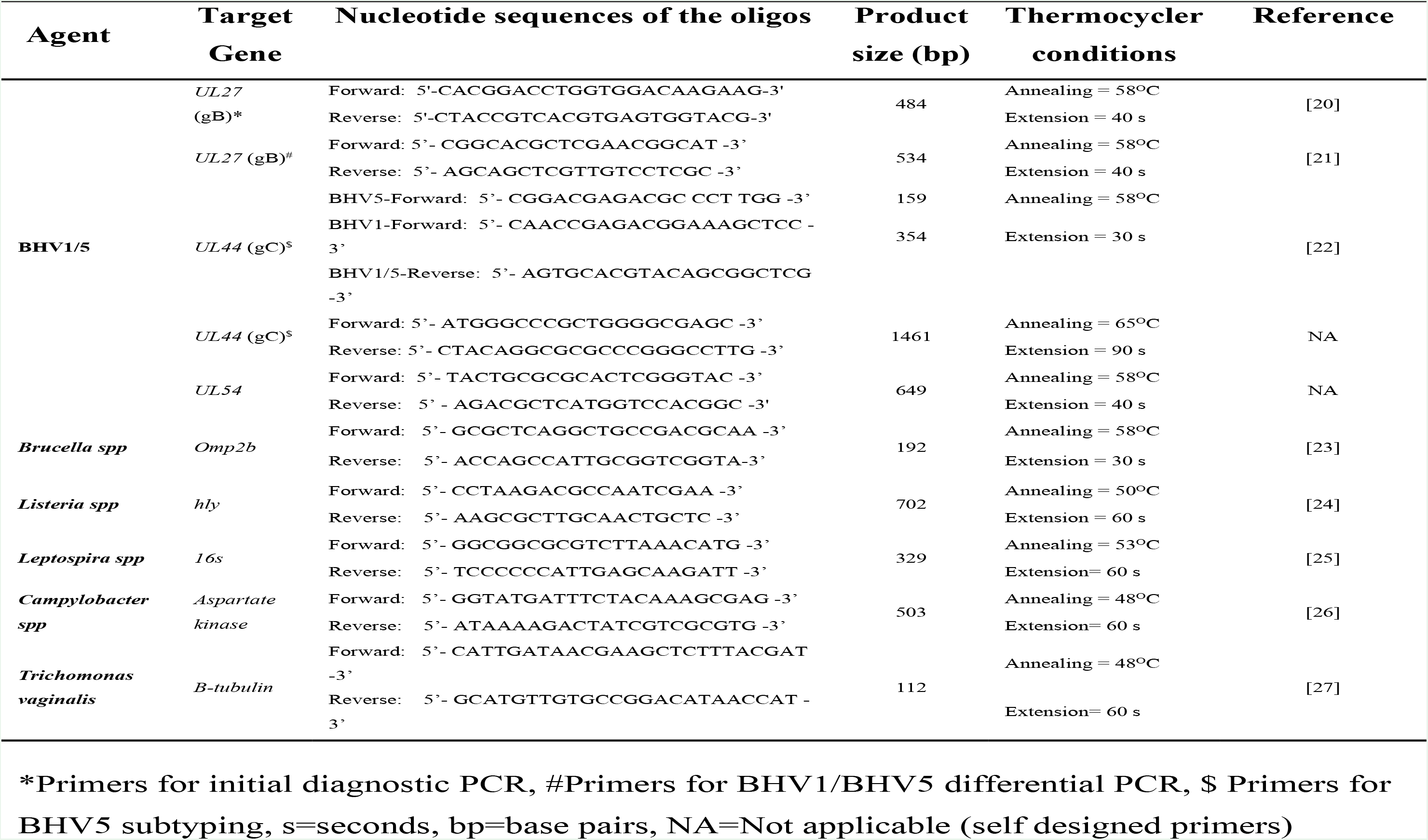
Primers used to amplify various agents involved in abortion.

### Virus isolation

For virus isolation, an aliquot of a biological specimen (500 μl filtrate) was used to infect a confluent monolayer of MDBK cells for 2 h followed by addition of fresh growth medium. The cells were observed daily for appearance of the cytopathic effect (CPE).

### Plaque assay

Plaque assay was performed as per previously described methods along with some modifications [28,29]. The confluent monolayers of MDBK cells, grown in 6 well tissue culture plates, were infected with 10-fold serial dilutions of virus for 1 h at 37°C, after which the infecting medium was replaced with an agar-overlay containing equal volume of 2X L-15 medium and 2% agar. The plaques were visible 5-6 days following infection (dpi). The agar-overlay was removed, and the plaques were stained by 1% crystal violet. The viral titres were measured in plaque forming unit/ml (pfu/ml).

### Virus neutralization assay

Virus neutralization assay was carried out to evaluate the antiviral antibodies in the infected animals. Serum samples were first heated at 56°C for 30 min to inactivate the complements. MDBK cells were grown in 96 well tissue culture plates at ~ 90 confluency. Two-fold serum dilutions (in 50 μl volume) were made in phosphate buffer saline (PBS) and incubated with equal volume of either BHV1 or BHV5 (10^3^ PFU/ml) for 1 h. Thereafter the virus-antibody mixtures, along with mock-infected control were used to infect MDBK cells. The cells were observed daily for appearance of CPE for determination of the antibody titres.

### Biotyping of BHV5

Based on the restriction endonuclease patterns, BHV5 has 3 subtypes, *viz*., A, B and C [22,30]. We subjected BHV5 to multiplex amplification of *UL27* and *UL54* genes by PCR followed by digestion by *BstE*II. The banding pattern generated was used to determine biotype of the BHV5 as described previously [22,30].

### Nucleotide sequencing

In order to further confirm the identity of the virus, *UL27, UL44* and *UL54* genes were amplified by PCR, gel purified using QIAquick Gel Extraction Kit (Qiagen, Hilden, Germany) and subjected to direct sequencing using both forward and reverse PCR primers. Duplicate samples were submitted for sequencing and high-quality sequences were deposited in the GenBank database with an Accession Numbers viz; MN852441 (*UL27*), MN852442 (*UL44*) and MN852443 (*UL54*).

### Phylogenetic Analysis

Nucleotide sequences from *UL27, UL44* and *UL54* genes (BHV5//India/2018) were edited to yield 447, 1368 and 585 bp fragments respectively using BioEdit version 7.0. These sequences, together with the representative nucleotide sequences of BHV1 and BHV5 available in the public domain (GenBank) were subjected to multiple sequence alignments using CLUSTALW (http://www.ebi.ac.uk/clustalw/index.html). Phylogenetic analyses were conducted using MEGA X. [31]. In order to evaluate the evolutionary history of the strain as well as phylogenetic relationship with different lineages, a Neighbor-Joining concatermeric phylogenetic tree comprising of *UL27, UL44* and *UL54* genes was generated. Test of phylogeny was performed using Maximum Composite Likelihood method and the confidence interval was estimated by a bootstrap algorithm applying 1,000 iterations. Molecular phylogeny and genetic relatedness of the isolated BHV5, with the rest of the BHV1/BHV5 strains were calculated using % similarity and pairwise distances.

## Results

### Identification of the agent(s)

For demonstration of the etiological agent in the aborted cattle, we extracted the DNA from the vaginal swabs and subjected it for amplification of BHV1, *Campylobacter spp, Brucella spp, Leptospira spp, Listeria spp and Trichomonas vaginalis* genome, the agents which are commonly associated with abortion in cattle. Among the bacterial/parasitic agents, amplification could not be observed except for *Brucella spp* (data not shown). Amplification of a 484 nt long PCR fragment with BHV1-specific primers primarily indicated the presence of BHV1 in vaginal swabs (**Fig. 1a**). Among the three aborted cattle, two were found positive for both BHV1 and *Brucella* whereas one was positive only for BHV1 genome. The PCR product (BHV1) was subsequently gel purified and submitted for direct sequencing. Nucleotide sequence similarity using NCBI BLAST revealed a close homology with BHV5 strains, rather than with BHV1 strains. This finding was intriguing because BHV5 had never been reported from India. When we re-examined the nucleotide sequences of the primers [32] that were used to amplify *UL27* gene by PCR, the sequences were found to be conserved among BHV1 and BHV5 genome. Therefore, the primer could amplify *UL27* gene of both BHV1 and BHV5.

**Fig. 1.**
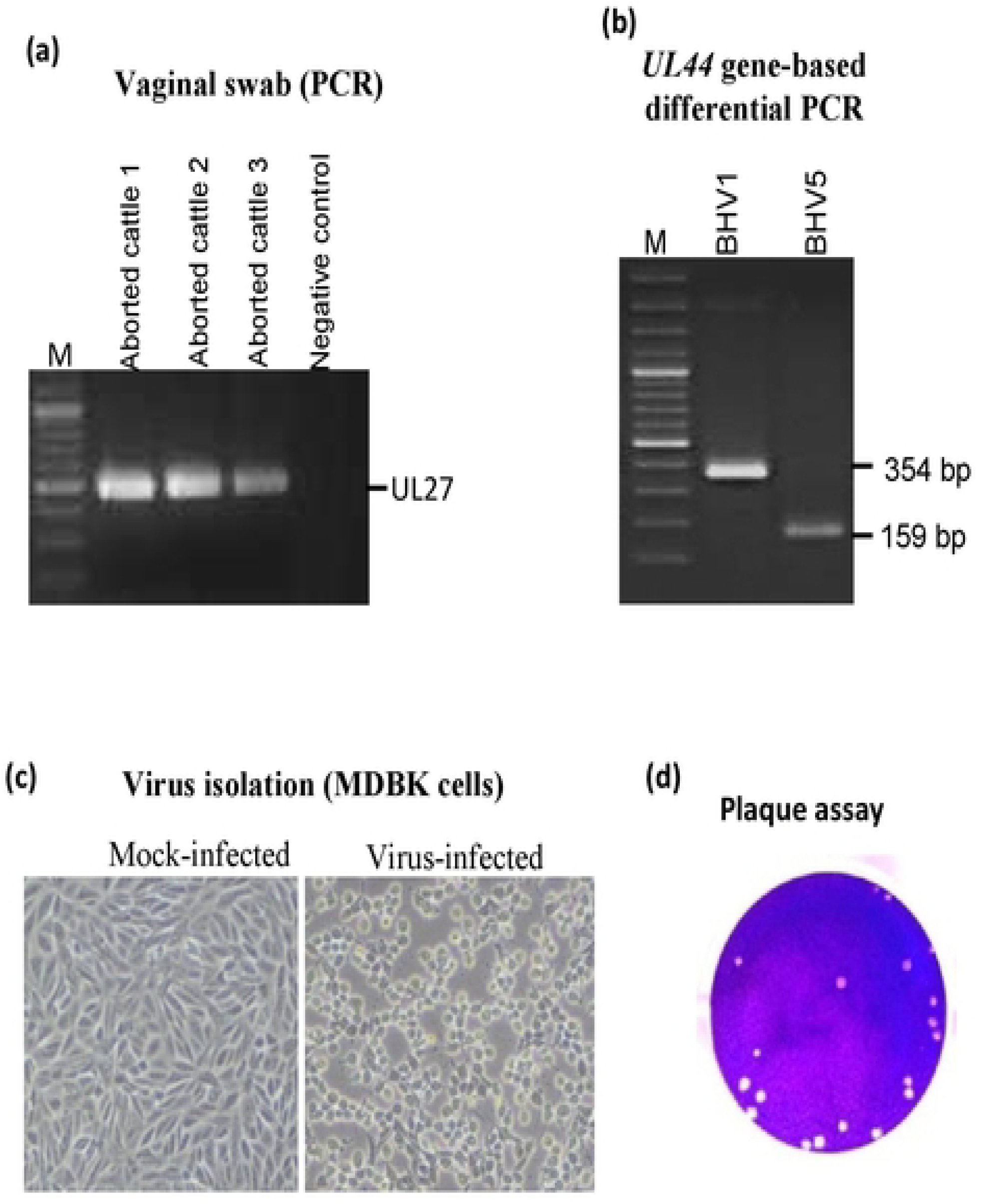
Isolation and identification of the agent. **(a) *Amplification of BHV1 genome in vaginal swab.*** Virus was recovered from the vaginal swabs in DMEM followed by DNA extraction and PCR to amplify *UL27* gene of BHV1. (b) ***BHV1/BHV5 differential PCR.*** PCR was carried out to amplify *UL44* as per the method described by Claus *et al*. PCR amplification of *UL44* from BHV1 (reference) and BHV5 (sample) resulted in amplification of 354 bp and 159 bp fragments respectively. (c). ***Virus isolation in MDBK cells.*** 500 μl of the virus recovered from the vaginal swab was used to infect MDBK cells. Cytopathic effect was observed at 3^rd^ blind passage. Virus-infected and mock-infected cells are shown. (d) ***Plaque assay.*** Confluent monolayers of MDBK cells were infected with 10-fold serial dilutions of the virus for 1 h at 37°C followed by replaced the medium with an agar-overlay. The plaques were visible at 5-6 days after infection. The agar-overlay was removed, and the plaques were stained by 1% crystal violet. The viral titers were measured in plaque forming unit/ml (pfu/ml).

The virus was also subjected to differential PCR as described previously [21]. As anticipated, PCR amplification of the *UL44* gene from BHV1 (reference) and BHV5 (sample) resulted in amplification of 354 bp and 159 bp fragments respectively **(Fig. 1b)** which further confirmed that the virus detected in vaginal swab was BHV5 and not BHV1.

### Virus isolation

The virus recovered from the vaginal swabs was used to infect MDBK cells, however no CPE was observed even up to 7 days post-infection. Thereafter, the infected cells were freeze-thawed twice and the resulting supernatant (called first blind passage) was used to reinfect fresh MDBK cells. In the third blind passage, the CPE characterized by cell rounding, ballooning and degeneration was observed at ~36 h post infection in virus-infected but not in mock-infected cells (**Fig. 1c**). Viral genome could be amplified in MDBK-infected cell culture supernatant at the 3^rd^ blind passage (data not shown). The isolated virus also formed plaques in MDBK cells which were visible within 5-6 days following infection **(Fig. 1d).** The virus infected cell culture supernatant had a titre of 1.2*10^7^ plaque forming unit per millilitre (PFU/ml) at the 3^rd^ blind passage. For bulk production, the MDBK cells were infected at MOI of 0.1 of BHV5 followed by virus harvest at 48 hpi by rapid freeze-thaw method. The bulk virus had a titre of 1.4*10^7^ PFU/ml. The virus was deposited in the microbial repository at NCVTC, Hisar bearing an Accession Number VTCCAVA218 and named as BHV5/*Bos taurus*-tc/India/2018/Bhilwara.

### Biotyping of BHV5

Based on the restriction endonuclease pattern, BHV5 has 3 subtypes, *viz*., A, B and C. By employing a previously described method, we also subjected the isolated BHV5 for subtyping [22]. As per the method, multiplex PCR amplification of BHV5 *UL27* and *UL54* genes **(Fig. 2a)** and their subsequent digestion by *BstE*II produces following banding patterns: (i) Type A: *UL27* (363 bp, 171 bp) and *UL54* (420 bp, 249 bp) (ii) Type B: *UL27* (534 bp) and *UL54* (420 bp, 249 bp) (iii) Type C: *UL27* (363 bp, 171 bp) and *UL54* (649 bp). In our study, PCR amplification of *UL27* and *UL54* genes and their subsequent digestion by *BstE*II produced four DNA fragments viz; 420 bp, 363 bp, 249 bp and 171 bp **(Fig. 2b)** which were suggestive of subtype A of BHV5/India/2018/Bhilwara.

**Fig. 2.**
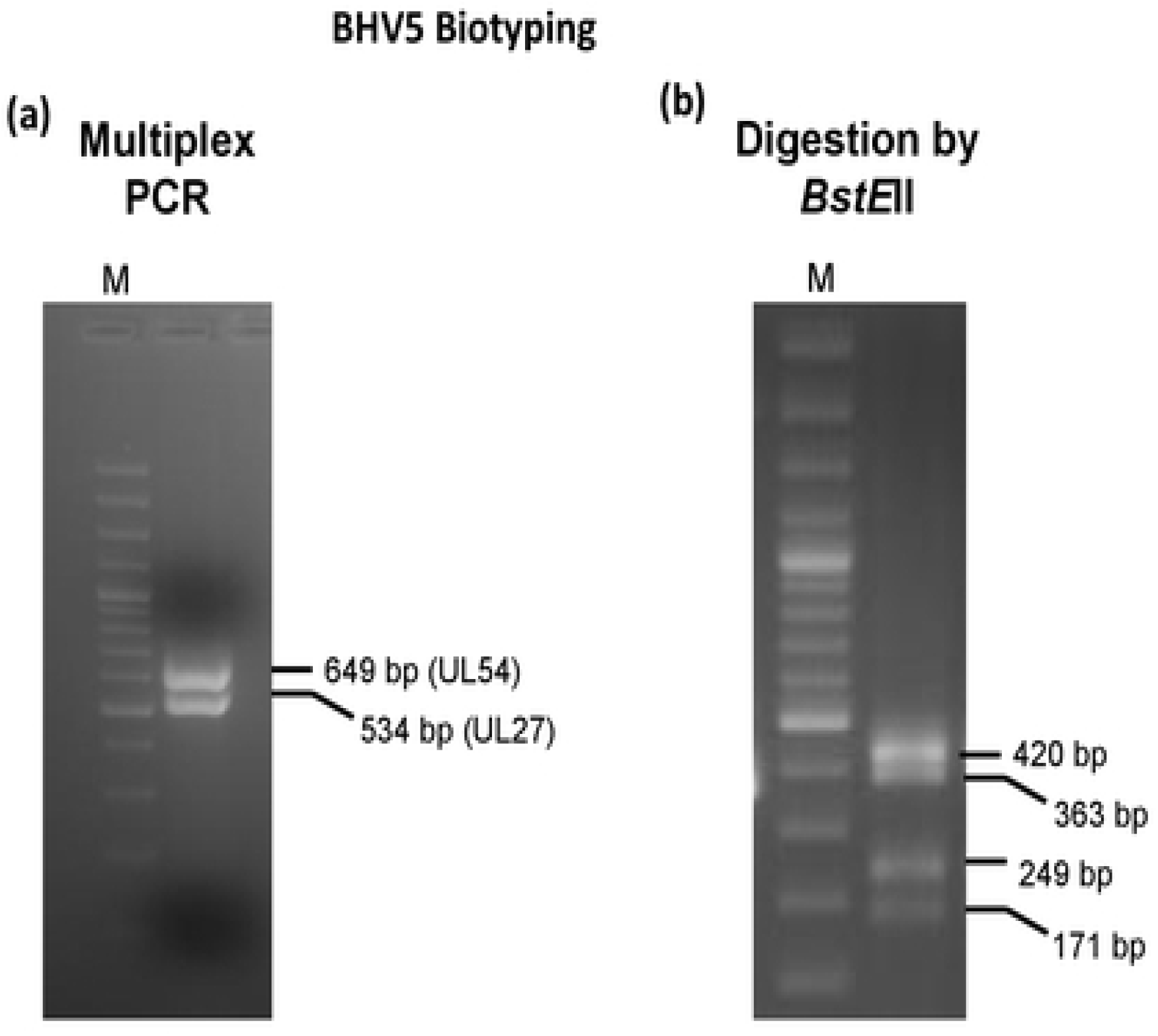
BHV5 subtyping: **(a)** Amplification of *UL27* and *UL54* genes of BHV5 by multiplex PCR resulted in amplification of 534 bp and 649 bp fragments respectively. **(b)** Digestion of PCR products (multiplex-*UL27* and *UL54* genes) by *BstE*II resulted in amplification of 420 bp, 363 bp, 249 bp and 171 bp fragments.

### Genetic relatedness

In UL44 (envelope glycoprotein C), the nucleotide (nt) and amino acid (aa) identities with other BHV5 strains were found to be in the range of 98.5-100% and 98.1-100% respectively and the highest identity (100%) was depicted with a Brazilian isolate (KY549446.1). The nt and aa identities in the same region with BHV1 isolates were in the range of 91.6-96.7% and 90.0-95.9% respectively. At the *UL54* region the nt identities ranged between 99.4-100% and the same were 90.4-99.5% in the partial glycoprotein B region (UL27) as compared to the other BHV5 isolates.

### Phylogenetic analysis

Neighbour-joining phylogenetic tree comprising of *UL27, UL44* and *UL54* genes was constructed to ascertain the evolutionary relationship of the virus with other BHV5 isolates in the public database. The Indian isolate clustered with the Brazilian BHV5 isolates with 100% bootstrap suggesting that the virus is closely related with BHV5 isolates from Brazil **(Fig. 3)**.

**Fig. 3.**
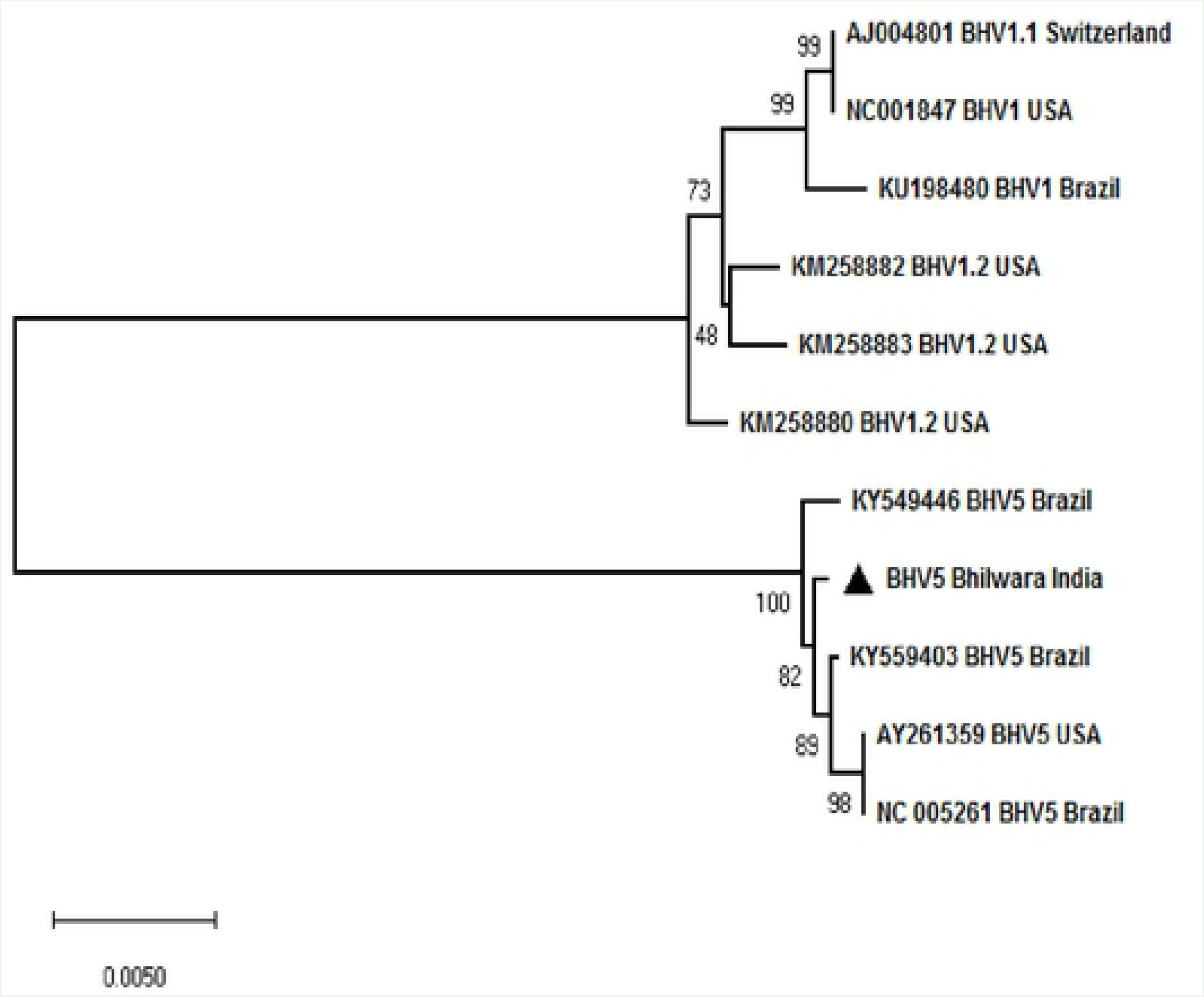
Phylogenetic analysis: Nucleotide sequences from *UL27, UL44* and *UL54* genes (BHV5//India/2018) were edited to 447, 1368 and 585 bp fragments respectively, using BioEdit version 7.0. These sequences, together with the representative nucleotide sequences of BHV1 and BHV5 available in the public domain (GenBank) were subjected for multiple sequence alignments Phylogenetic analyses were conducted using MEGA X. To evaluate the evolutionary history of the strain as well as the phylogenetic relationship with different lineages, a concatemeric Neighbor-Joining method tree was generated. Test of phylogeny was performed using Maximum Composite Likelihood method and the confidence intervals were estimated by a bootstrap algorithm applying 1,000 iterations. The tree is drawn to scale, with branch lengths in the same units as those of the evolutionary distances used to infer the phylogenetic tree.

### Analysis of recombination

To elucidate any evidence of recombination between the BHV5 isolates, by employing RDP4 programme, we also carried out recombination analysis in the *UL44* gene with the available sequences in the public database. The default parameters available in RDP4 programme *viz*., RDP, GENECONV, BOOTSCAN, MAXCHI, Chimera, SISCAN and TOPAL were employed to identify the recombination breakpoints and the parental strains. However, the analysis did not reveal any evidence of recombination (data not shown).

### Detection of antiviral antibodies

We also evaluated the levels of antiviral antibodies in three aborted and three apparently healthy cattle belonging to the same farm. An antibody titre of 8-128 was observed irrespective of the abortion history **(Table 2)** suggesting an active BHV5 infection in the herd. Serum samples of all the aborted cattle were also positive for antibodies against *Brucella* **(Table 2)**.

**Table 2:**
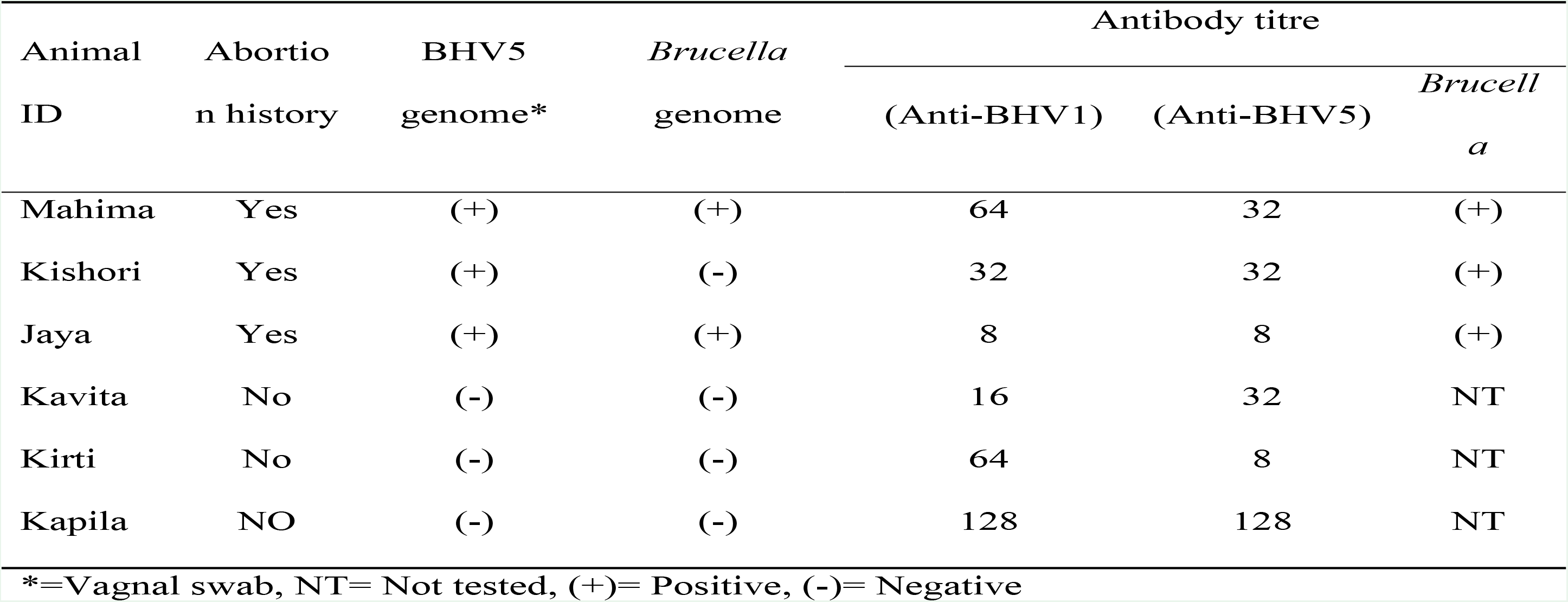
Detection of viral genome and antiviral antibodies in aborted and apparently health animals.

| Animal ID | Abortion history | BHV5 genome* | *Brucella* genome | Antibody titre |  |  |
| --- | --- | --- | --- | --- | --- | --- |
|  |  |  |  | (Anti-BHV1) | (Anti-BHV5) | *Brucella*<br>a |
| Mahima | Yes | (+) | (+) | 64 | 32 | (+) |
| Kishori | Yes | (+) | (-) | 32 | 32 | (+) |
| Jaya | Yes | (+) | (+) | 8 | 8 | (+) |
| Kavita | No | (-) | (-) | 16 | 32 | NT |
| Kirti | No | (-) | (-) | 64 | 8 | NT |
| Kapila | NO | (-) | (-) | 128 | 128 | NT |

\*=Vaginal swab, NT= Not tested, (+)= Positive, (-)= Negative

## Discussion

BHV5 distribution is restricted to South American countries, particularly Argentina [9], Uruguay [10] and Brazil [11]. Only a few cases of this disease have been reported from other countries [12,13,15,16]. BHV5 has never been reported from India. Clinical findings, detection of antiviral antibodies, virus isolation, BHV1/BHV5 differential PCR, biotyping and sequence and phylogenetic analysis of *UL27, UL44* and *UL54* genes confirmed the association of BHV5 subtype A in the aborted cattle. To the best of our knowledge this is the first report on the presence of BHV5 infection in India.

DNA extracted from vaginal swabs of aborted cattle showed the presence of BHV1 (three aborted cattle) and *Brucella* (2 aborted cattle). Other agents such as *Campylobacter spp*, *Listeria spp*, *Leptospira spp*, *Trichomonas vaginalis* were not detected. Rapid generation of CPE (within 36 hrs) in MDBK cells, a characteristic of BHV1 further indicated association of BHV1 in aborted cattle. However, nucleotide sequences revealed a close homology with the BHV5 strains, rather than with the BHV1 strains. When the corresponding nucleotide sequences of the primers [32] used to amplify *UL27* gene were re-examined, they were identical in BHV1 and BHV5 genomes. Therefore, the primer could amplify *UL27* gene of both BHV1 and BHV5 strains.

In order to further confirm, the isolated virus (BHV5//India/2018/Bhilwara) and a reference BHV1 control were subjected to differential PCR as described previously (Claus et al., 2005). As anticipated, PCR amplification of the *UL44* gene from BHV1 and BHV5 resulted in amplification of 354 bp and 159 bp fragments, respectively, which further confirmed the identity of the virus as BHV5, not BHV1.

Based on the restriction endonuclease patterns [22,30], BHV5 has 3 subtypes, *viz*.,; A, B and C. Type strains for subtypes A, B and non-A-non-B, are the Australian strain N569, the Argentinean strain A663 and Brazilian strains, respectively. The banding patterns generated following PCR amplification of *UL54* and *UL27* genes (BHV5//India/2018/Bhilwara) and their subsequent digestion by *BstE*II were clearly suggestive of BHV5 subtype A.

BHV5 infection in cattle usually causes meninogoencephalitis [3], although few reports also suggest the involvement of the reproductive tract [4]. Besides demonstration of the virus (BHV5) and BHV5-specific antibodies in the aborted animals, no obvious neurological signs could be recorded in any of the aborted cattle. There are reports suggesting interspecific recombination between BHV1 and BHV5 [33]. However our analysis did not reveal any evidence of recombination. Besides, we could not detect evidence of BHV1 infection also in the farm. Although reproducing clinical BHV5 disease under experimental conditions is a tedious task, some laboratories have developed a rabbit model of encephalitits [34–38]. Although the isolated virus in our study was ~99% identical with the Brazilian BHV5 strains, its complete genetic characterization (whole genome sequencing) as well as its ability to produces encephalitis in natural host and/or in rabbits needs to be elucidated which is beyond the scope of this manuscript. Likewise, precise role of the isolated virus (BHV5/India/2018/Bhilwara) and/or other infectious agents (coinfection) in inducing abortion in the cattle farm needs to be determined.

Besides demonstration of the etiological agent(s), serum samples were also found positive for both anti-BHV1/5 and anti-Brucella antibodies suggesting an active BHV1 and *Brucella* coinfection in the herd. Some serological evidences have shown BHV1 and *Brucella* coinfection [39–42], however, demonstration of agent (BHV1/5 and *Brucella*) has been a rare event. It is a matter of further study whether *Brucella* and BHV5 coinfection have synergistic or antagonistic effect on disease severity [43].

BHV1 vaccine induces cross-protection against BHV5 disease in cattle [7]. Naturally occurring or vaccine-induced anti-BHV1 antibodies are believed to reduce the occurrence of BHV5-associated disease in BHV1 endemic areas [44]. However, with reasons precisely unknown, this does not apply to the epidemiology and transmission of BHV5 in South America. Even though Argentina and Brazil have a high percentage of BHV1 seropositive cattle (24.8–84.1% and 19–85%, respectively) [reviewed in reference [3]], both countries have reported several cases of BHV5 associated meningoencephalitis. Furthermore, serological tests which can differentiate anti-BHV1 and anti-BHV5 antibodies are not available. Thus, the actual prevalence of BHV5 infection and hence economic significance remains unknown. Furthermore, the phenomenon of viral interference between BHV1 and BHV5 needs to be studied.

Like Europe and USA, since India has no specific programme for the detection and identification of BHV5 infected animals, it might have been overlooked despite its presence. The concerned farmer always practiced natural service and never performed artificial insemination for breeding purposes. However, the bulls were procured from nearby state (Surendranagar, Gujrat). India import semen from several other countries including Brazil, therefore the possibility of introduction of BHV5 via semen from Brazil and/or other countries cannot be ruled out.

Very few studies have been undertaken on the development of BHV5 vaccine, firstly because of its limited geographical distribution and secondly reproducing the clinical BHV5 disease under experimental conditions is difficult [3,45]. Countries with frequent BHV5 outbreaks along with high prevalence of BHV1 have successfully employed BHV1 vaccines for protection against BHV5 infection [7,8]. However, the levels of anti-BHV5 antibodies induced by BHV1 vaccine is usually low and of shorter duration [46]. Therefore, each BHV1 vaccine should be carefully tested for potential cross-protection against BHV5.

To conclude, we provide a strong evidence of BHV5 infection in Indian cattle for the first time. The isolated virus would be useful for developing diagnostic, prophylactic and therapeutic agents to combat BHV5 disease in India. The finding may necessitate inclusion of BHV5 test protocol in testing of semen for sexually transmitted diseases.

## Acknowledgements

This work was supported by the Science and Engineering Research Board, Department of Science and Technology, Government of India, Grant number CRG/2018/004747 to N-Ku and Indian Council of Agricultural Research, Grant Number IXX11882 to N-Ku and S-B. The funders had no role in study design, data collection and analysis, decision to publish, or preparation of the manuscript.

## Author contribution

N-Ku and R-T designed the experiments. N-Ku, Y-C, N-Kh, R-K and H-K performed the experiments. N-Ku, S-B, R-T and B-NT wrote the manuscript.

